# Brain Injury Localization in Electromagnetic Imaging using Symmetric Crossing Lines Method

**DOI:** 10.1101/2021.09.22.461428

**Authors:** Guohun Zhu, Alina Bialkowski, Stuart Crozier, Lei Guo, Phong Nguyen, Anthony Stancombe, Amin Abbosh

## Abstract

To avoid death or disability, patients with brain injury should undertake a diagnosis at the earliest time and accept frequent monitoring after starting any medical intervention. This paper presents a novel approach to localize brain injury using the intersection of pairs of signals from symmetrical antennas based on the hypothesis that healthy brains are approximately symmetric that the bleeding targets will lead to significantly different amplitude and phase changes if one of pair of transmit signals cross targets. The scattered signals (S-parameters) are acquired using 100 realistic brain models and 150 experimental data measurements. Firstly, three pair of horizontal antennas are used to detect target crossing which line and in which hemisphere in low frequency bands and estimate the size using high frequency bands. Then, an intersection of two pairs of antennas are identified the position of the target. Finally, a heat map is used to visualise the stroke brain. The results indicate that crossing pairs of antenna signals from the hemisphere with a blood mass exhibit significantly different signal amplitude in the graph features compared to those without the target (p<0.003). The experiments show that our novel localization algorithm can achieve an accuracy of 0.85±0.08 Dice similarity coefficient based on 150 experimental measurements using an elliptical container, which is suitable for brain injury localization.

## I. Introduction

**E**LECTROMAGNETIC (EM) imaging technologies have been attracting widespread interest from researchers around the world due to their potential as a portable, low-cost and non-ionizing medical diagnostic system [1]–[5]. This technique has recently been implemented for different medical applications, such as the detection of stroke [1], [3], breast cancer [5], [6] and skin cancer [7]. Two important applications are the detection of breast cancer [8] or traumatic brain injury (TBI) at an early stage. TBI is a major cause of mortality and disability. Intracranial haemorrhage (ICH) is the most important complication of TBI. A mass lesion refers to an area of localized brain injury that must be removed at the earliest time possible for reducing the mortality rate. In most scenarios, people suffering from TBI are not administered timely treatment because the symptoms, such as decreased level of consciousness, cannot always be observed in the early stages. Clinical experiences indicate that the size and location of ICH are critical for clinicians in an operation, therefore a fast and accurate method for ICH localization and size estimation is of prominent importance.

Currently, most studies based on EM imaging aim to identify the types of tumors, stroke or intracranial bleedings (IBs) [1], [2], [8]–[11]. The state-of-the-art methods for brain injury localization can be categorized into three types: radar-based, tomography, and machine learning based methods. The radar-based methods and relevant results were reported in [4], [12], [13]. Tomography-based methods are capable of mapping the dielectric properties of brain tissues, including the abnormal ones, hence providing the localization of brain bleedings [3], [14]. In general, these methods need to solve complex inverse problem to identify the target from the solved dielectric distribution [15]. Furthermore, it was indicated in [16] that when certain a-priori information was used, such as the patient’s magnetic resonance imaging (MRI) scan, results from tomography approaches could obtain 0.24 relative error on 35 subjects. Machine learning-based methods [2], [17]–[19] can avoid the computationally expensive solving of the inverse problem. However, these methods are more suitable for stroke sub-type classification [2], [18]. Apart from stroke classification, the studies in [17], [19] also attempted to localize the stroke position using support vector machines (SVMs) or the artificial fish swarm algorithm. Unfortunately, both methods require a huge amount of data for training even with a single phantom or simulated brain model. These requirements are usually difficult to satisfy in clinical scenarios [2]. In fact, while these three approaches have their own merits, they also have drawbacks. For example, the tomography-based methods are ill-posed and may not converge to the correct solution, in addition to taking long computational time. The radar-based methods often fail to correctly localize and shape the target, particularly in the case of heterogeneous tissues [20]. EM imaging using machine learning methods is still not accurate in localizing brain injury without large amounts of training data. These limitations restrict practical utilization of portable electromagnetic systems, especially in emergency applications. In addition, all these methods require precise and permanent positioning of the antennas [21]. This requirement could degrade the performance of these methods during practical applications.

This paper presents a novel symmetrical crossing line algorithm (SCLA) to localize the position and predict the size of a TBI. This is based on the hypothesis that a healthy brain is nearly symmetric with respect to the left-right axes [22], [23]. The assumption gives rise to the potential that anomalies can be detected based on the significant difference in symmetric pairs of signals across this axis. The inputs to the algorithm are the S-parameters collected from an antenna array surrounding the head as well as the approximate positions of the antenna ports. The array used to test the technique is composed of 16 antennas which are approximately equally spaced around the head. only transmitted signal are utilized as the transmitters, whereas all reflected signals are ignored. To avoid the influence from amplitude variations, the signal measured from each antenna is converted into a weighted graph. Electromagnetic waves transmitted between two antennas whose signal path pass through bleeding area will have a significantly different weight of the graph features compared with signals in the mirrored antenna pair in the other hemisphere. Based on this idea, the IB can be localized by detecting the intersection of two transmitting lines with the highest weighting coefficient. A size constructing algorithm (SCA) helps to estimate the target centre and the size from three crossing lines. In total, 100 simulated data and 150 experimental data are used to evaluate the algorithm.

## II. Experimental Data and Methodology

Fig 1 shows the work process adopted in this study, which can be summarized as follows:

**Fig. 1:**
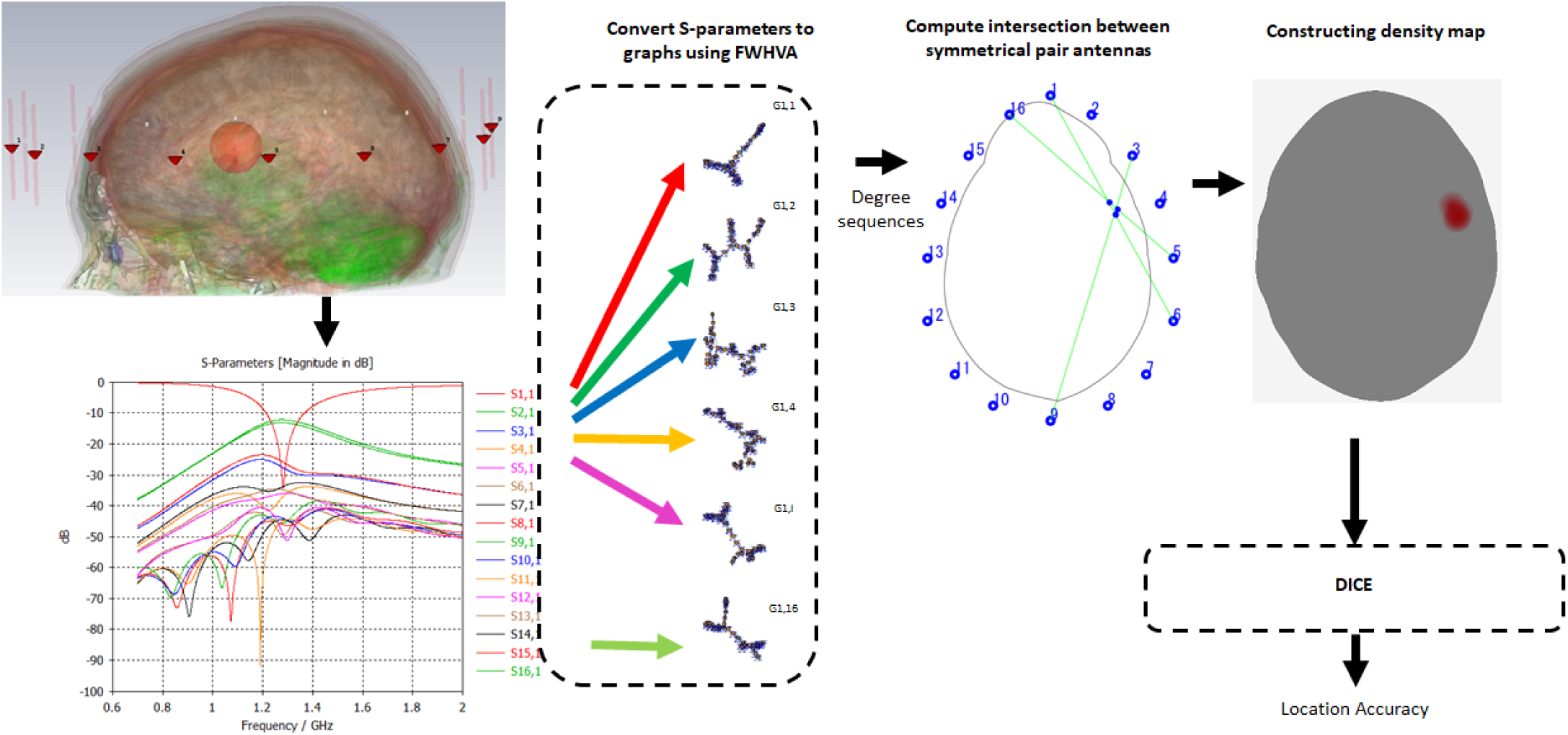
Flowchart of the proposed localization procedure.

- A patient is monitored by an EM imaging device with *n* antennas after a suspected traumatic brain injury
- The scattered data (*S*_*i,j*_, where *i, j* ∈ [1 … *n*]) is collected by an antenna array
- An inverse FFT is applied to convert these signals to *n × n* time series signals, and each time series is transformed into a graph using the Fast Weighted Horizontal Visibility Algorithm (FWHVA) [24].
- Each symmetric pair of signals is compared to determine mismatches, which are then used to localize the bleeding based on the intersection of the direct signal paths:
  – The centre of the bleed is obtained by a symmetric crossing line algorithm (SCLA).
  – The size of the target is estimated using a size constructing algorithm (SCA) based on the SCLA output.
- The Dice similarity coefficient (DICE) is computed to calculate the localization accuracy.

### A. Brain models and simulation set-up

This study investigates the proposed SCLA algorithm by evaluating the simulation results of eight realistic brain models. The brain models used in this study include detailed Zubal [25] and Duke [26] phantoms that were derived from multiple imaging modalities. In addition, 6 other brain models (named P103111 to P111716), which were built from segmented MRI images [27], are utilized. Each model includes eight tissues (blood, skin, skull, CSF, grey matter, white matter, cerebellum, and ventricles), The details of the phantoms are listed in Table I. The number of voxels indicate the dimensions of the phantoms e.g. 280 × 330 × 280 shows that the X, Y slices are 280 × 330 pixels, and the 280 indicates the number of slices in the Z-direction.

**TABLE I:** Details of three type of models

| Phantom IDs | Number of tissues | Resolution (mm) | Number of voxels |
| --- | --- | --- | --- |
| Zubal [25] | 58 | $1.1 \times 1.1 \times 1.4$ | $256 \times 256 \times 128$ |
| Duke [26] | 52 | $1.0 \times 1.0 \times 1.0$ | $610 \times 310 \times 161$ |
| P103111-P111716 | 8 | $0.7 \times 0.7 \times 0.7$ | $280 \times 330 \times 280$ |

The proposed SCLA algorithm is evaluated using the electromagnetic simulator, CST Microwave Studio. To build phantoms with intracranial bleeding from the healthy models of Table I, each healthy model was injected with an elliptical shape target (blood), which emulates a TBI, for which each axis was randomly assigned a length of 1 mm to 10 mm and the position was randomly distributed in the X and Y plane limited from 76 to 152 for Zubal and 76 to 218 for others respectively. Z-direction slices ranging from 100 to 230 were used.

### B. Hypothesis

The hypothesis of this paper is that a brain containing an intracranial bleed will have an imbalance between symmetric pairs of signals, based on the assumption that a healthy brain is approximately symmetric [22], [23]. As electromagnetic waves pass through a domain, the signals are altered in amplitude and phase owing to different dielectric properties of the media that the signals pass through. Thus, when signals arrive at two antennas with the same distances, the amplitudes or phases will be significantly different if only one of them passes through a high contrast bleeding. In additional, when size of IB is large, both high frequency and lower frequency have the same trends compared with left or right hemisphere. Conversely, only high frequency has the significant changes if the size of IB is small.

Firstly, we define a differential opposite pairs (DOP) and differential neighbour pairs (DNP) associated with two antennas as equation 1 and equation 2, respectively.

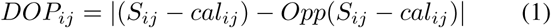

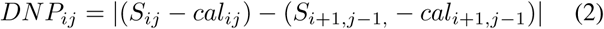

where *S*_*ij*_ is the scatter parameters between pairs antennas *i* and *j* and Opp() indicates the symmetrically opposite antenna pair (mirror pair), originating from the same antenna *i*. where *cal*_*ij*_ is the features from the scattered signals measured with an elliptical homogeneous isotropic media in both simulation and experiment, without a head present. If the head is exactly layout in the centre of antenna arrays, the *cal* can be removed from equations 1 and 2. Certainly, regarding the value from low frequency (0.5-1.35 GHz), the symbol is labeled as *LDOP* and the *HDOP* is from high frequency bound (1.35-2 GHz).

#### Lemma 2.1

Given three pairs of symmetry antennas *DOP*_1,8_, *DOP*_2,7_ and *DOP*_3,6_, then the pair crossing the targets has obvious notches in between 0.8 − 1.3GHz, and the corresponding time series has a slower decay.

Fig. 2 (a) shows a 16-antenna array with antennas positioned symmetrically around the left-right axis of the head. The locations of these antennas are provided in supplementary Table 1.

**Fig. 2:**
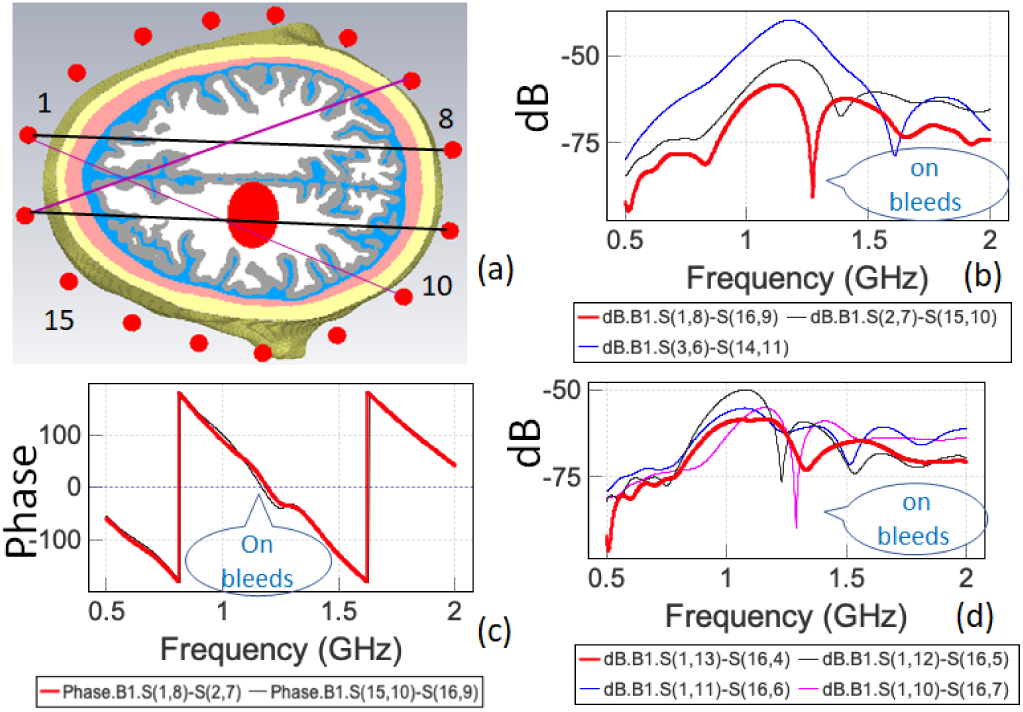
(a) A brain surrounded by a 16-element electromagnetic-imaging antenna array showing the direct signal paths between antenna pairs where the blood mass is indicated as a red ellipsoid inside the brain. (b) three differential S-scatter coefficients pairs, where *S*(1, 8) − *S*(16, 9) has a notch (c) (15, 10) − (16, 9) has larger phase about 1G (d) S(1-10)-S(16,7) has a notch.

Fig. 2 (b) shows the S-parameters corresponding to the transmit coefficients of the three antenna pairs in Lemma 2.1. It is clearly that S(1,8)-S(16,9) has an obvious notch about 1.25GHz with the smallest value(−90DB). Therefore, the target is on the pair of (1, 8) or (16, 9), but not other two pairs.

#### Lemma 2.2

Given two of adjacent pairs antennas in the same hemisphere, *DNP* (1, 8), and *DNP* (16, 9) or *DNP* (2, 7) and *DNP* (15, 10), then the pair crossing the targets has obvious larger phase about 1*GHz* but will have smaller phase about 1.25G than the other hemisphere.

Fig. 2 (c) shows the phase on S(15,10)-S(16,9) has a larger phase about 1GHz than those hemisphere pairs *S*(2, 7) − *S*(1, 8).

#### Lemma 2.3

Given two of adjacent pairs antennas in the left and right hemisphere, from *DOP* (1, 10), *DOP* (1, 11), … and *DOP* (1, 13), then the pair crossing the targets has an obvious notch between 1 − 1.3*GHz*.

Fig. 2 (d) shows the phase on S(1,10)-S(16,7) has a deep notch 1GHz than those hemisphere pairs *S*(2, 7) − *S*(1, 8). Because the lemma 2.2 shows that the target is on the side of pair of 16, 9. Thus, the target was located by the pairs of (1,10) and (16,9). Certainly, it is possible to use antenna 8 and 9 to find another lines which cross the targets using the Lemma 2.4

#### Lemma 2.4

Given two of adjacent pairs antennas in the left and right hemisphere, from *DOP* (8, 12), *DOP* (8, 13), … and *DOP* (8, 15), then the pair crossing the targets has an obvious notch between 1 − 1.3*GHz*.

However, the Δ*S* method with the minimum DB might introduce errors in other antenna arrays because the transmit coefficient have multiple notches (such as ultra wide band antennas). In addition, it is difficult to detect the weakly differences in the S-parameter, such as Fig 2(c). Thus, this paper introduces an oblique visibility graph approach to detect the abnormal pair transmit lines and align the signals.

### C. Normalizing electromagnetic signals using graph features

Regarding the example brain in Fig. 2, the signals from different pair antennas have a different amplitude and phase. More importantly, different head sizes also affect these parameters. To normalize the amplitude and align the different phases, an oblique visibility graph (OVG) [24] is employed to convert those phase or amplitude sequence into a graph. OVG have been widely used in time series, such as EEG [28] and ECG [29]. This paper applies it to frequency domain processing by processing the amplitude and phase, respectively..

From a mathematical point of view, an amplitude or phase sequence of a signal can be regarded as a time series, *X*_*j*_(*j* = 1, …, *t*), which is mapped to a graph *G*(*V, E*) by the following steps: a data point *x*_*j*_ is mapped into a node *v*_*j*_ in *G*; an edge *e*_*jk*_ between (*v*_*j*_, *v*_*k*_) from two points (*x*_*j*_; *x*_*k*_) exists if and only if:

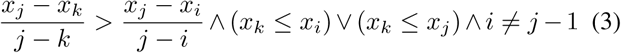

Fig. 3 shows how signals from a real system between 11 ns to 15 ns are converted into an oblique visibility graph (OVG). In Fig. 3 (a), a portion of the phase signal received from Fig. 2 (c). In this plot, the 5^*th*^ point can horizontally see the 12^*th*^ point but cannot horizontally see the 13^*th*^ point because the 12^*th*^ point blocks the visibility between 5^*th*^ point and the latter. It can be seen that the signals decay rapidly in the time domain as shown in Fig. 3 (a). However, HVG can effectively detect the sine wave and abnormal signals.

**Fig. 3:**
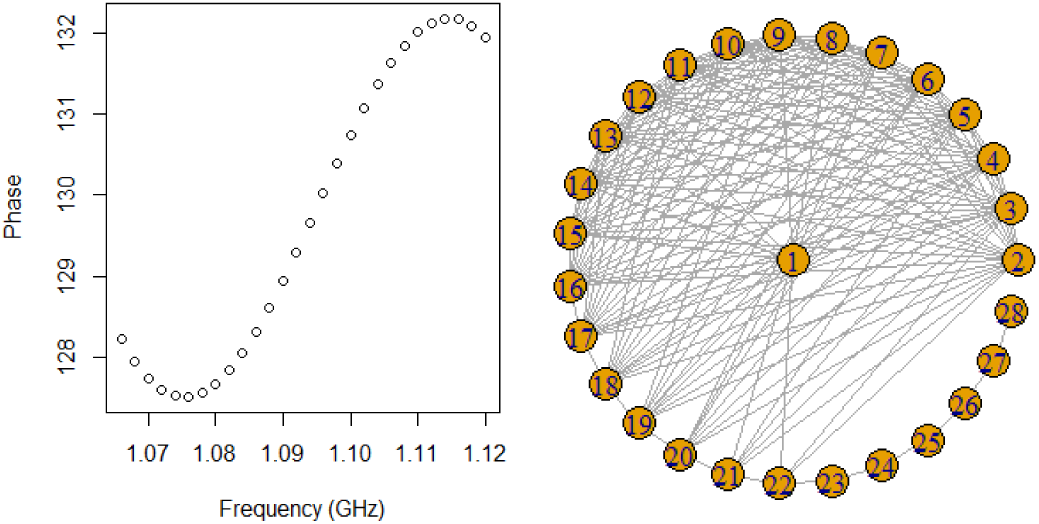
(a) 28 points of phase signals from 1.07G Hz to 1.12G Hz, and (b) An OVG mapped from the signal in (a).

In a complex network such as an oblique visibility graph (HVG), node degree and degree sequence are two basic characteristics which can be used to describe the graph. The degree *d*(*v*_*i*_) of node *v*_*i*_ is the number of connected edges from *v*_*i*_. As shown in Fig 3(b), the node *d*(*v*_28_) = 1 and *d*(*v*_27_) = 2. It is notices that the notch of the wave will have a large degree *d* which can indicator the notch places, such as nodes 2, 3, 4, 5 and son on.

Fig 4 illustrate two OVGs from two phase signals of antenna pairs *S*(2, 11) − *S*(15, 6) and *S*(2, 12) − *S*(15, 5). It can confirm the hypothesis of Section II.B that the transmit line that crosses targets will introduce a lager phase and graph degree changes compared with the opposite transmit line. Certainly, the phase changes is weakly but the graph degree changes are significantly.

**Fig. 4:**
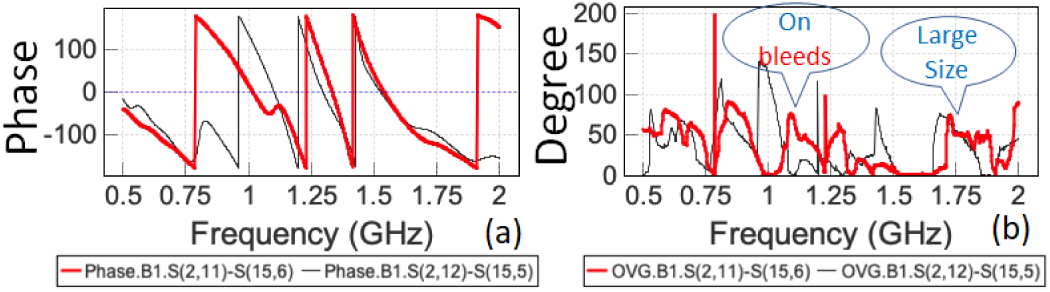
(a) phase signals from DOP(2,11) and DOP(2,12) of Fig 2 and (b) Two OVGs mapped from the signal in (a) where crossing lines from target can be easy to marked with a larger degree about 1.1-1.2 Ghz and the size might be identified in high frequency about 1.75Ghz.

## III. SCLA Algorithm

Based on the hypothesis that a brain containing intracranial bleeding will have a significant difference in the measured signals of symmetric pairs of antennas, we can locate the bleeding based on which pairs exhibit large differences.

The SCLA algorithm is divided into four steps: mapping the magnitude and phase series into graphs, identifying the correct quadrant of the bleeding location as in the left or right hemisphere, picking candidate intersection of pairs lines, and constructing the heat maps. The first step is based on OVG algorithms. The next three steps are described in the following sections.

### A. Localizing the hemisphere of the bleeding targets

In clinical cases, it is also important to detect which hemisphere (left/right) the bleeding lesion occurs. Although experts might detect it based on clinical symptoms. A naïve approach to detect whether the blood mass is in the left or right hemisphere is by comparing the features (such as amplitude, mean degree or strength) from the left signal to those of the right. However, this method can rarely provide an accurate answer due to the slight asymmetry of the head. This section presents a simple method to detect whether the bleeding is in the left or right hemisphere based on Lemmas 2.1 and 2.2.

#### Algorithm 1 Intersection algorithm

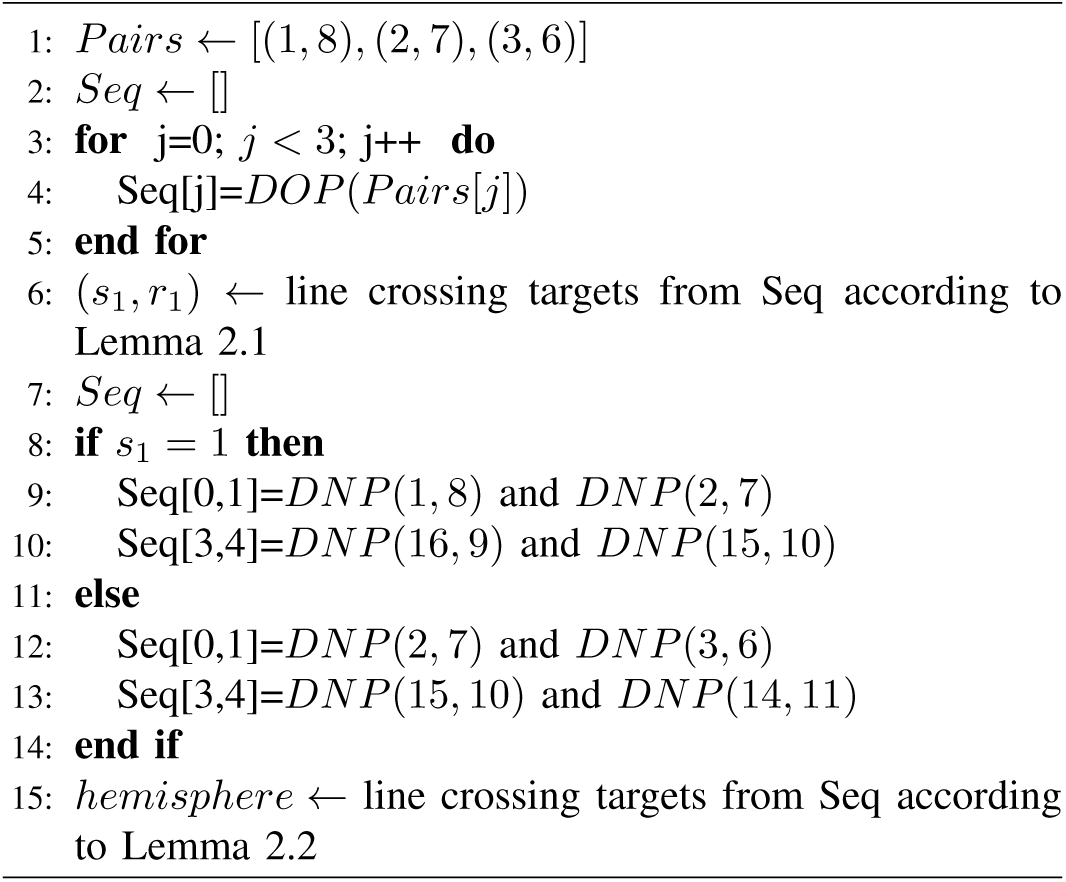

One example of the algorithm is from the Fig 2. The (*s*_1_, *r*_1_) = (1, 8) based on Fig 2. Then *DNP* (1, 8) will be compared with the opposite pairs *DNP* (16, 9) as shown on Fig 2, where the *DNP* (16, 9) has larger degree about 1.25G Hz position. Thus, the target is on the left side pair line (16,9).

### B. Picking the intersection of an antenna pair and its mirror

Algorithm 1 only obtains a horizontal line (*s*_1_, *r*_1_) closest to the target. This section will provide an algorithm that two oblique lines that crossing the targets.

#### Algorithm 2 Intersection algorithm

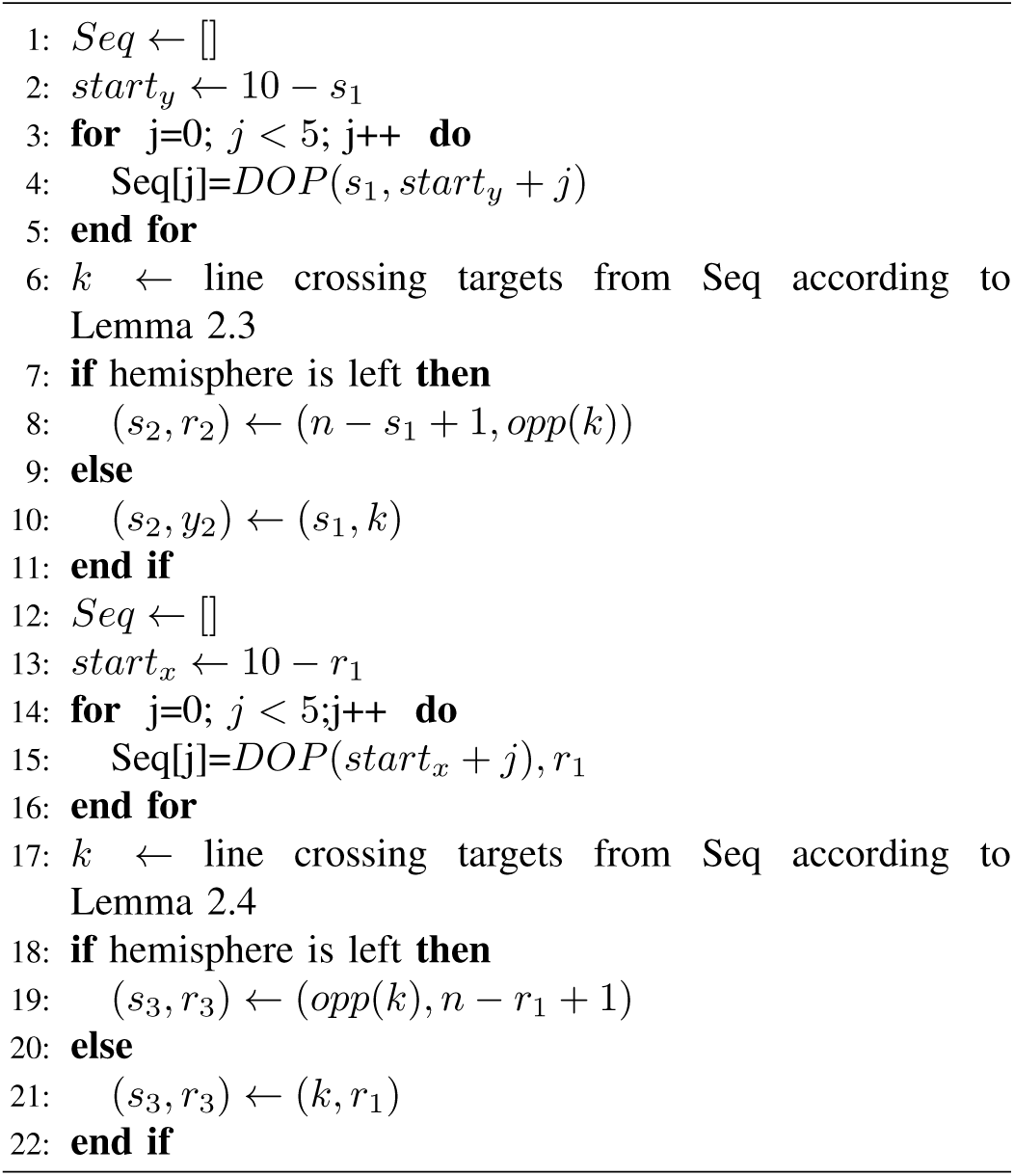

Algorithm 2 produces two intersecting lines *p*_2_ and *p*_3_ from (*s*_2_, *r*_2_) and (*s*_3_, *r*_3_). As shown in Fig 2, the output of algorithm 1 are (1, 8) and left hemisphere, Then the output by Algorithm 2 becomes (1, *r*_2_) and (8, *r*_3_). Conversely, if it is right hemisphere, then the output of Algorithm 2 is (16, *y*_1_) and (9, *y*_2_). Then, the intersection co-ordinates (*p*_2_, *p*_3_) can be calculated based on (4) and (5) when four antenna positions (*x*_1_,*y*_1_), (*x*_2_,*y*_2_), (*x*_3_,*y*_3_) and (*x*_4_,*y*_4_) are provided.

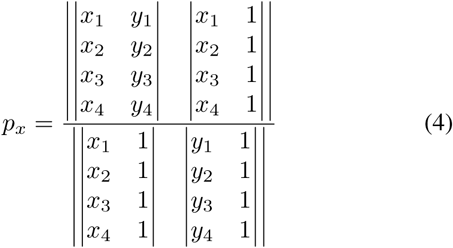

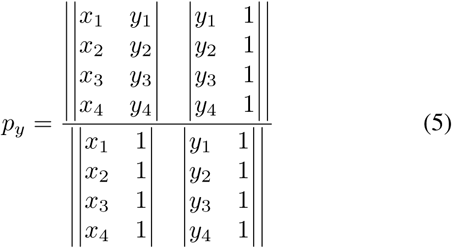

### C. Heat map constructing algorithm

This section describes a constructing heat map algorithm to obtain the size and the shape of the bleeding from the intersection of lines pairs from section III-B. The algorithm uses intersected heat mapped shape as a lesion, which requires the output of Algorithms 1 and 2 as input values, such as I=*{R*_0_, *R*_1_, *R*_2_*}*. The pseudo code is listed below.

a. Calculate the maximum distance between the intersections I from the SCLA algorithm. For example, there are three intersections *R*_0_ = (*x*_0_, *y*_0_), *R*_1_ = (*x*_1_, *y*_1_), and *R*_2_ = (*x*_2_, *y*_2_), the maximum distance *md* = |*R*0 − *R*1|.
b. Detect the maximum degree on high frequency (about 1.7G Hz) of the output pair (*s*_1_, *r*_1_) of Algorithm 1, if there has larger degree on the *NOP* (*s*_1_, *r*_1_), then heat radius is assigned as 20*mm* otherwise 40*mm*.
c. Draw each intersection point with the assigned heat radius.

### D. DICE

To evaluate the performance of localization, the Dice similarity coefficient (DICE) is used to calculate the overlap of the estimated size of the IB (*S*_*t*_) with the real IB area (*S*_*d*_). The DICE, also called the overlap index, is the most used metric in MRI segmentation. It is defined by

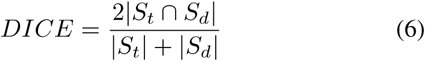

The DICE can be viewed as a similarity measure over two bleeding sets. If two image contains the same area coordinate points, the DICE is equal to 1.0, while if two bleeding sets have no overlap, it is equal to 0.0.

### E. Statistical analysis of head rotation

To investigate the difference between objects with targets and those without targets, a quantitative statistical test was performed on the two sets of graph features. A p-value< 0.05 was obtained, demonstrating that there is a significant difference in the graph features of objects with vs without targets.

## IV. Results on Simulated Data

The algorithms are implemented in *C* and *R* languages. Eight phantoms were employed for testing. Each phantom is constructed with five different bleeding observations by injecting an elliptical shaped blood mass with the dielectric properties of blood. In total, 28 brain model observations were used for evaluating the algorithm performance in simulations. In addition, a simple phantom with a small glass cylinder filled with distilled water to emulate a TBI is used to verify the algorithm with a 14 antenna array in experiments. The glass cylinder is moved clockwise to two different distances per antenna, and the S-parameter data is collected twice to measure the anti-noise performance of SCLA. Thus, a total of 56 simulated data are collected and tested. The mean DICE value of 94 testing samples is 0.78 and the maximum DICE value is 0.92.

### A. Test the algorithm when rotating antenna

To test the tolerance of the algorithm to the antenna positions, this subsection evaluates the antenna Z-rotation tolerance in the simulation experiment. Rotating the antenna array along the Z-axis will significantly affect the algorithm performance because the other two dimensions (X and Y axis) can be easily aligned by ears and eyebrows. In experiments with healthy volunteers, we found that subjects were biased to left or right on the Z-axis but few of them rotated their heads on the X or Y-axis.

Figures 5 shows an observation from a brain model from the seven tissue brain models. The red target of Figures 5(a) indicates a bleed target on the ground truth, where the figures 5(b) shows the SCLA algorithms’ output.

**Fig. 5:**
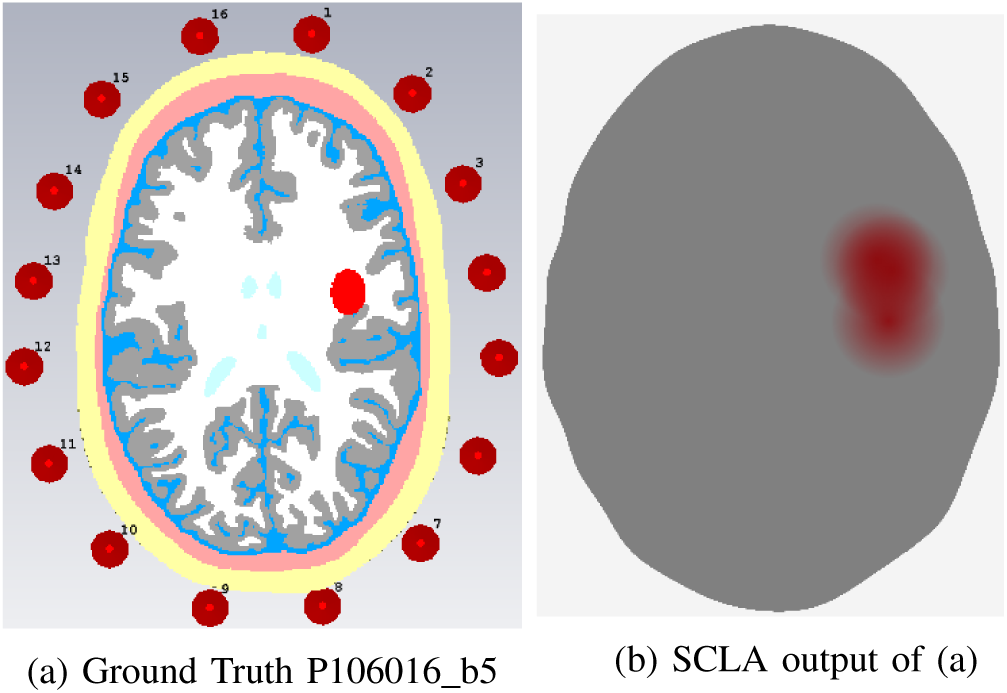
A test from model P106016 b5, where the ground truth positions with tissues are shown on the left and the SCLA algorithm result is shown without tissues on the right.

Fig 6 shows the same brain model as Fig 5 but the antenna array has three degrees of rotation in Figure 6. The Ground Truth of a Duke Model DICE between the SCLA output and the ground truth is approximately 0.08 in Fig 6, while the DICE is about 0.84 in the symmetrical case (Fig 5). Thus, this algorithm can tolerate a two-degree antenna rotating in all directions.

**Fig. 6:**
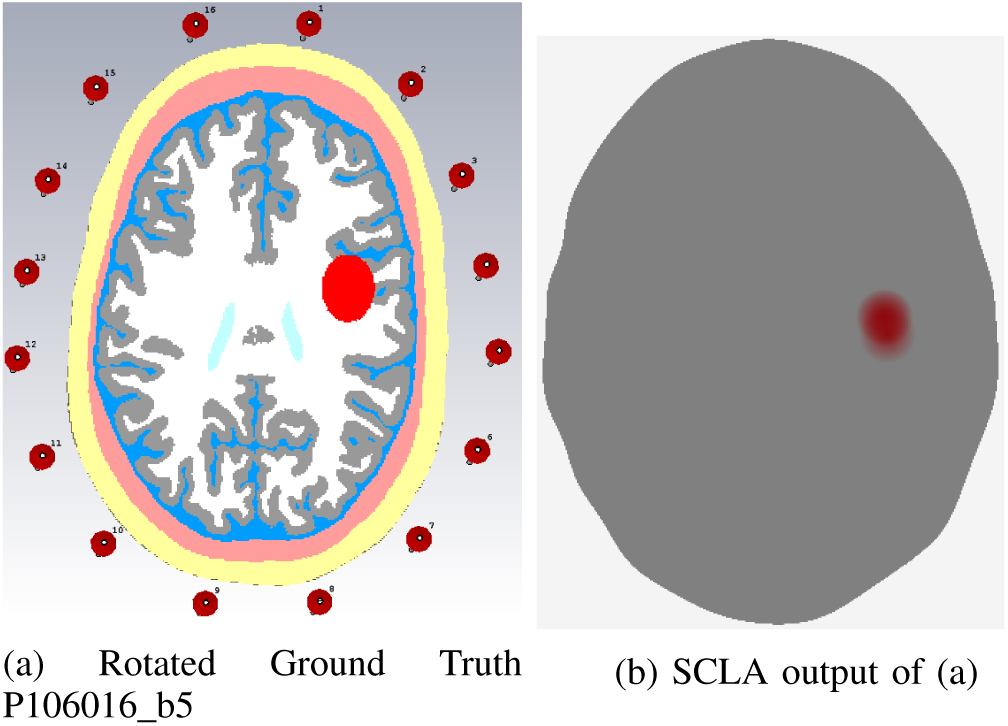
A test from model P106016 b5 is the same as Fig 5 but all antenna positions are rotated 3 degrees, where the ground truth positions with tissues are shown on the left and the SCLA algorithm result is shown without tissue on the right.

### B. Testing different antenna positions

Figures 7 and 8 show two observations from two brain models with intracranial bleeding (Duke and Zubal brain models, respectively) with different distances between the head and antennas. The green line indicates the direct signal path from antennas 1, 5, and 9. The crossing points are plotted in blue, whereas the estimated targets are drawn in red.

**Fig. 7:**
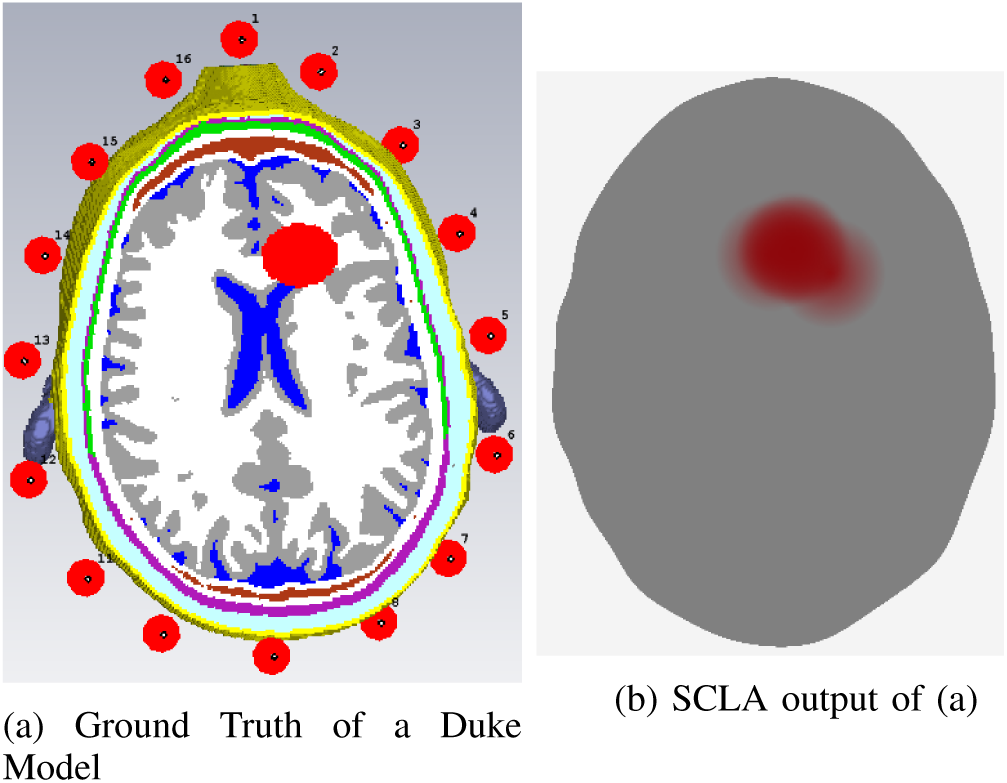
A test result on the Duke brain model. The ground truth positions with tissues are shown on the left and the SCLA algorithm result is shown without tissues on the right.

**Fig. 8:**
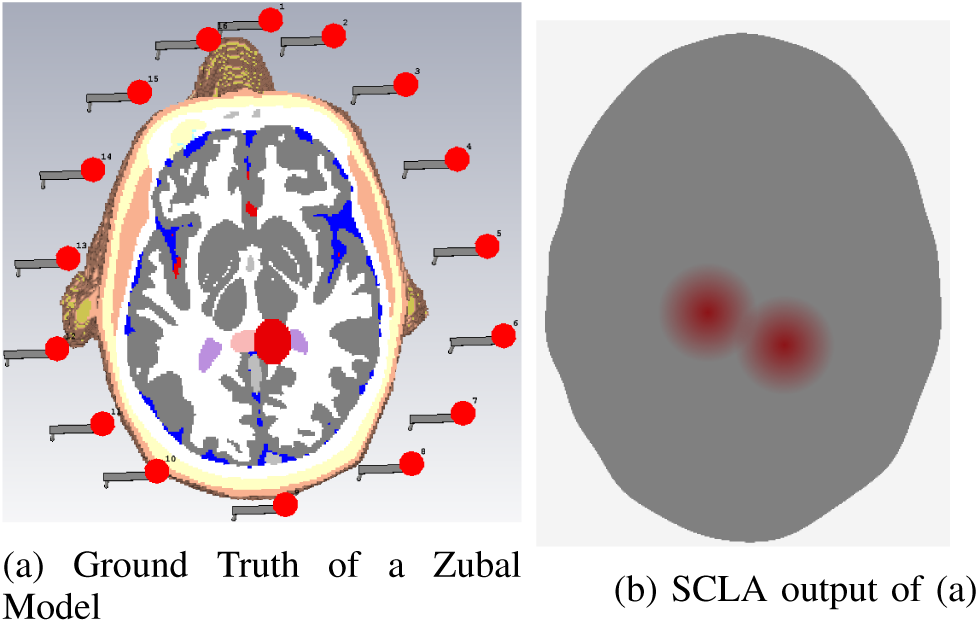
A test result of the Zubal brain model. The ground truth positions with tissues are shown on the left and the SCLA algorithm result is shown without tissues on the right.

It can be seen that the Duke and Zubal brain models are significantly different. In addition, the gap between head boundary and antennas in Fig 7 is smaller than those in Fig 8. Our SCLA was able to obtain the approximate location of the targets even though the two antennas have different placement and the distances between the antennas and the head boundary are also different.

## V. Results on experimental Data

### A. Experiments data

The SCLA was applied to locate simple targets in two experiments. One is a dielectric-loaded 14-waveguide antenna array system. Each waveguide antenna is filled with a coupling medium with a dielectric permittivity of 33 and a conductivity of 0.1. An elliptical container filled with 60:40 glycerol water mixture is used to mimic a head, and a water-filled glass cylinder with an outer diameter of 22 mm and wall thickness of 1 mm is used as a target. The second is a 16 Log-Periodic array system. The experimental device is shown in Fig 9(a and b).

**Fig. 9:**
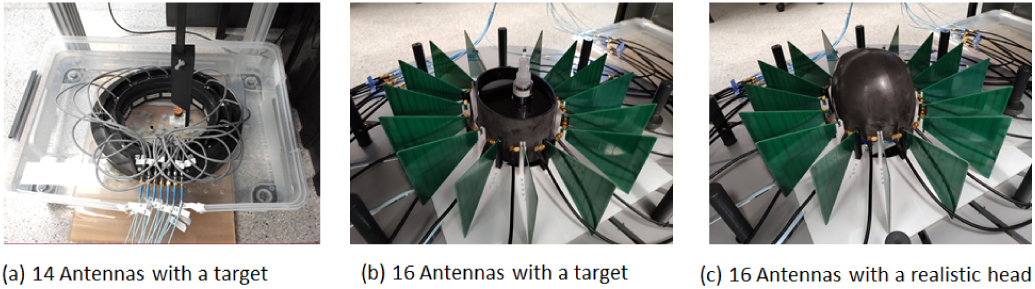
Three experimental platforms with 14 and 16 antennas respectively.

The target in Fig 9(a) is moved clockwise in front of each antenna with 10 mm and 14 mm distances, respectively. The S-parameter data was collected twice. Thus, a total of 56 targets were measured. A Keysight M9037A VNA was used to measure the signals, and a frequency scan from 0.5 GHz to 2 GHz with 2 MHz step size was used to provide a total of 751 frequency points.

The second experiment evaluated the algorithm performance using a CNC machine with a EM head scanner. There are 16 antenna array put on a Skin-Skull Bucket (SSB) as shown in Fig 10(a). The SSB was placed within the array imaging domain upside-down and filled with liquid which has average brain emulating dielectric property. It was placed on a centring mount within the array, to hold the SSB in the centre of the antennas. The pressure of the array membrane was then increased to 1 PSI. Following this, a solid haemorrhage emulating target (Fig 10(c)) was measured inside the SSB. These targets were moved along a 10mm spaced grid pattern in X and Y, with a fixed Z as low as possible within the bucket. In total, there were 94 points measured inside the SSB.

**Fig. 10:**
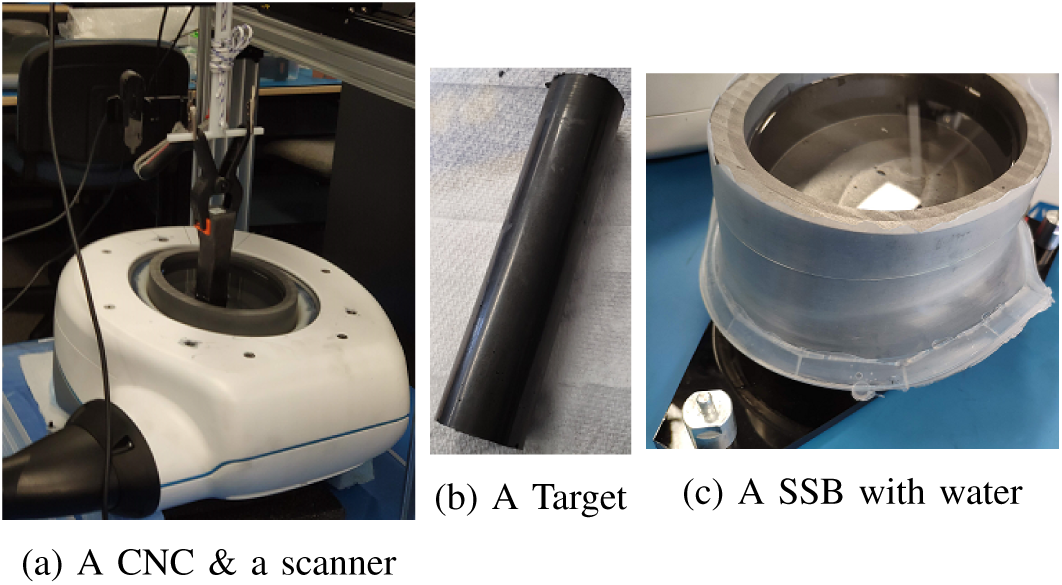
The SCLA output of two realistic phantoms.

For all these realistic systems, the measurement time is approximately 1.5 seconds per measurement on 16 antennas. The system uses a transmit power of approximately +10 dBm (10 mW), which is less than that used in mobile phones. Compared with an existing study which performed experiments at only a single frequency point [30]. Our system is slower due to a large number of frequency points (751 steps).

### B. Locating the target in a realistic container

As introduced in Section V-A, an elliptic container filled with distilled water was used to test a target which is a glass cylinder with an outer diameter of 22 mm and wall thickness of 1 mm, filled with distilled water. During each measurement, a CNC machine moves the target to one of 28 positions, and the distance between the between the target and the antenna aperture is 10 mm and 14 mm respectively. A movie was made in the Supporting files to illustrate the target movements. Fig 11 shows a SCLA output of a measurement, where the container with target was used for calibration in equation 1. The SCLA can correctly localize the target according to the collected S-parameters. The average DICE score for these 56 measurements was 0.85 *±* 0.08.

**Fig. 11:**
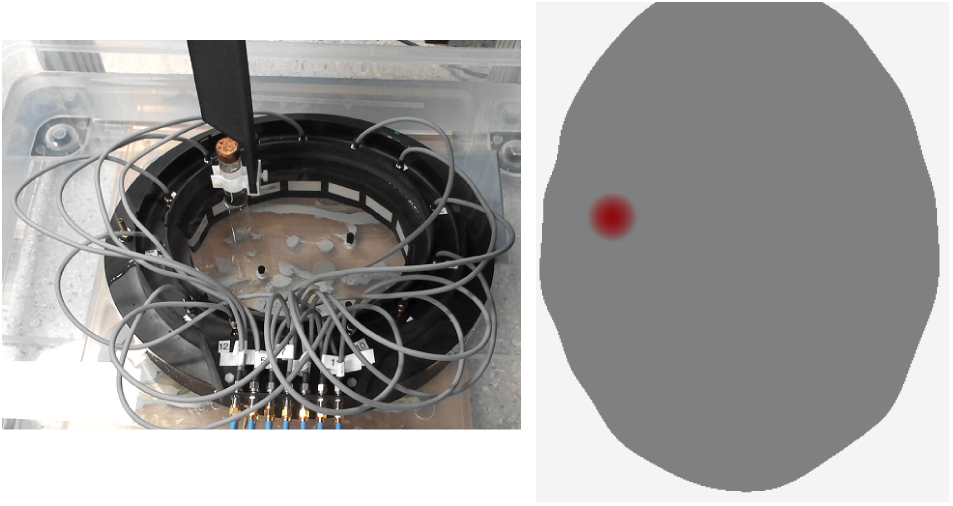
The experimental setup. The glass cylinder filled with distilled water was moved into a stable position near antenna 11 in a water filled container on the left and the SCLA algorithm result is shown in red on the right.

### C. Locating targets in asymmetric head phantoms

This section test the algorithms using 94 ICH targets which is placed in 94 positions inside a SSB container. One of the original position is shown in Fig 12 (a). It is clearly that the symmetric pair 4 − 12 and 5 − 13 can divide those targets into two groups as shown in Fig 12(b). For example, the 87^*th*^ target is 100% at the back and the 23^*st*^ target is 100% on the front. Then, the other pairs such as pairs 1 − 9 and 16 − 8 could divide the target into left or right hemisphere.

**Fig. 12:**
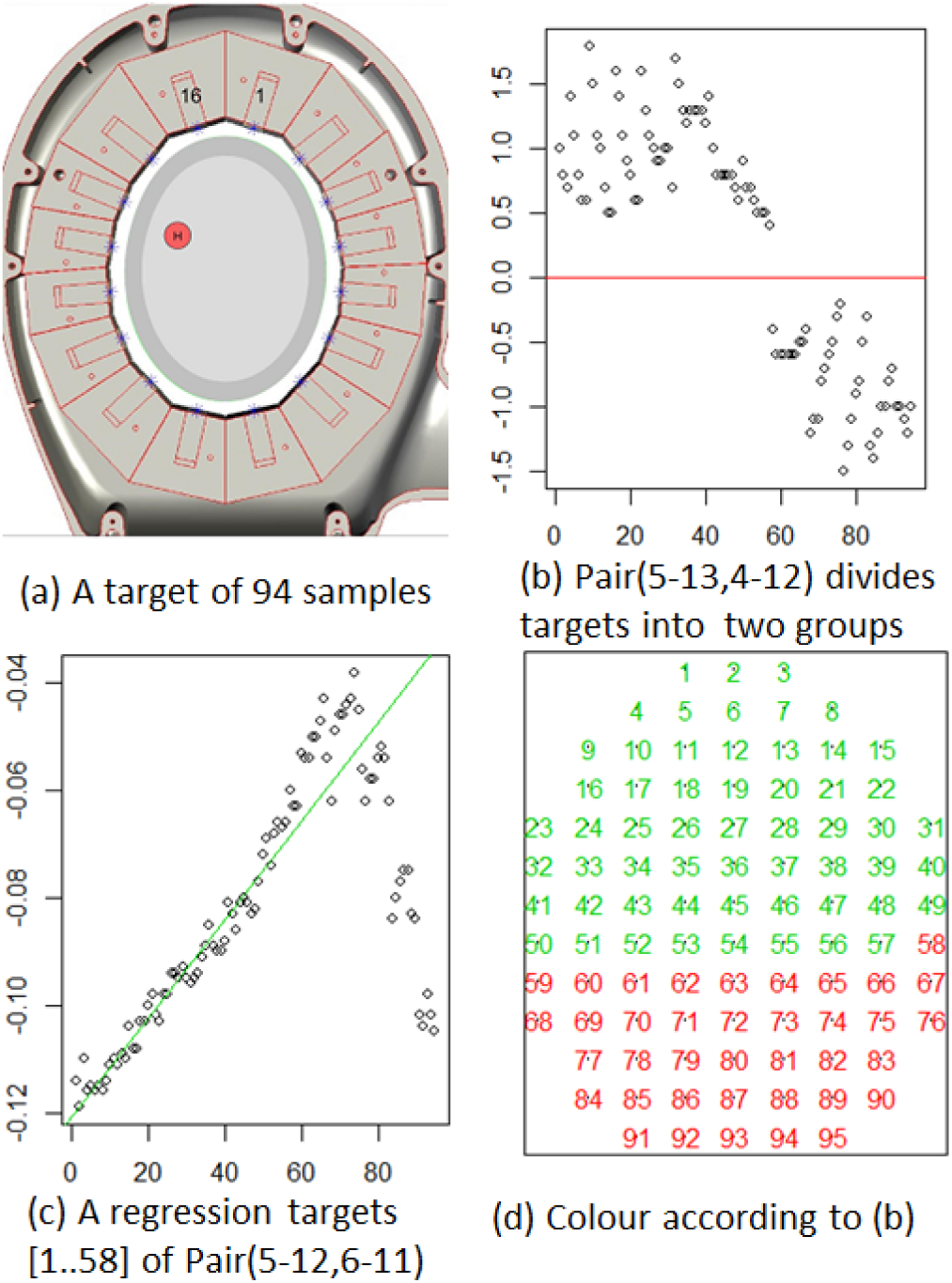
Two pairs antennas analysis of 94 points in a SSB container

It is easy to find that the measurement results support that the real head is not completely symmetrical. Fig 12 (c) shows that the differential degree features from neighbour pair 5 − 12 and 6 − 11. The value increase when target is near the transmit line between pair 5 − 12 and 6 − 11, such as targets 60 or 75. On the other hand, the position of target 1 − 58 has a linear regression relationship with those of *DNP*_5,6_. In that case, it is easy to obtain the target 23^*r*^*d* position by using the regression in Fig 12 (c) as we could detect the target is really in front area by detecting the *DOP*_4,5_ as shown in Fig 12 (d). Certainly, any target in back area could have the similar linear relationship with the pair of 4 *−* 13 and 3 *−* 14.

## VI. Discussion

In this study, we presented a novel symmetrical crossing line algorithm (SCLA) which only requires the antenna positions and measured signals to localize intracranial bleeding in the brain. The physical principle is that, compared to signals passing through two similar areas (non-target areas), a pair of symmetrical transmitted signals passing through the targets (bleeding) will generate significantly different amplitude and phase changes. Unlike conventional imaging algorithms, the output of the SCLA is the coordinates of the target instead of an image. Compared to other methods which need human or machine learning technology to provide the real coordinates from the medical image, our algorithm directly provides the real coordinates.

In fact, the SCLA might open a new direction for electromagnetic imaging which relies on the intersection of the direct signal paths between antenna pairs. Compared with radar-based detection methods, such as confocal imaging [31], the SCLA algorithm can obtain the location without knowing the boundary of the head and without estimating the average permittivity of the imaging subject. In fact, the SCLA algorithm is only based on the intersection of the two signal paths, which can obtain high localization accuracy even when the head boundary is unknown.

Compared with using multi-core CPUs [8] or using GPUs tomography-based methods, the SCLA method has much lower computational cost (less than 1 minute on an Intel Core i7-2600 CPU @ 3.40 GHz with 16G RAM), making it more suitable for use in emergency scenarios. In addition, the tomography-based methods are usually influenced by the selection of the antennas. For example, in [3], [32], antennas with *TM*_*z*_ polarization needed to be used to satisfy the integral equations, and in [33], directive waveguide antennas were used to restrict the radiation pattern so that the scattering problem could be approximated as a 2-D problem.

Compared with the machine learning-based methods which require a huge training data set [17], SCLA is based only on the on-site measurement and it is not a learning-based method thereby the substantial training data are not required.

Also, human heads have different sizes and shapes. Many existing algorithms fail to give an accurate image when the brain sizes and shapes are different. However, our SCLA algorithm is not affected by the boundary, size or shape. The only requirement is the positions of the antennas which is easy to obtain in a clinical situation. More importantly, the container and the target were filled with the same distilled water in Fig 11. Despite this, our proposed algorithm is still able to efficiently detect the changes caused by the 1 mm thick glass cylinder only.

The hypothesis of this method is assuming that the healthy human head is approximately symmetric across the major and minor axes (i.e., across the sagittal plane, separating the left-right of the brain, and across the central coronal plane, separating the front and back of the brain). Symmetric pairs of antennas across these lines of symmetry should have similar signals within a healthy head. If symmetric pairs of antennas measure different signals, this indicates an abnormality such as bleeding, and this observation can be used to facilitate localisation. In our realistic system, each antenna is just 10 mW and the measurement is about 1.5 ns. There are 8 transmitting antennas in total. This is very safe because it is only the equivalent to 0.01s of a mobile phone transmission time.

Sections IV and V demonstrate that SLCA can obtain cerebral hemorrhage images from experimental phantoms and simulation data. However, two reasons could lead to difficulty in measuring the DICE in experimental data compared with those of simulated cases. Firstly, it is that the CNC can introduce error during movements, while the simulation model has none of this type of error. Secondly, shifts or rotations of the phantom or the human head might introduce errors during the measurement as shown in section IV-A. Currently, there is no effective tool to detect whether the phantom or head is placed in the center of the antenna array. It can only rely on manual and visual inspection to observe whether the head or phantom is rotated or shifted. Therefore, it is possible to obtain low or zero DICE in real experiments despite high DICE being achieved in simulation in the same situation. To overcome this limitation and to acquire a higher DICE score, we can use multiple measurements to determine whether the crossing line of the algorithm are from the same pair of antennas. In the future, a method to automatically detect rotation or shifts of a head or phantom will be developed.

Currently, the algorithm does not calculate the electrical properties of the bleeds. However, it can identify target localization using differential degree values between two antennas. Thus, it is possible to measure a target in 3-dimensional imaging by moving the system up or down on the z-axis using a 3-dimensional calibration. Each scan is a slice, and then all slices could be merged into a 3-dimensional case. In addition, it is possible to detect other abnormal tissues, such as tumors or ischemic in the future.

## VII. Conclusion

Using a novel symmetrical crossing line algorithm, this paper is the first to localize an intracranial bleeding mass without using radar, tomography or machine learning methods. Based on the hypothesis that healthy brains are approximately symmetric, the algorithm is able to accurately localize and predict the shape of bleeding masses that cause a non-symmetric response in the electromagnetic signals. The algorithm was evaluated on 150 measurements with head phantoms of varying size, shape and resolution, and antenna types, and was able to accurately locate bleeding masses with an average Dice score of 0.85. Compared with existing electromagnetic imaging and localization algorithms, our approach is less sensitive to changes in head size, coupling mediums and antenna positions, directly outputs the coordinates of the bleeding mass, and does not require any training data. It is also significantly faster, localising the bleeding mass position within one minute. The speed and accuracy of the method shows great potential for use in emergency clinical applications.

